# ß2-Arrestin germline knockout does not attenuate opioid respiratory depression

**DOI:** 10.1101/2020.08.28.272575

**Authors:** Iris Bachmutsky, Adelae Durand, Kevin Yackle

**Affiliations:** Deparment of Physiology, University of California-San Francisco, San Francisco, CA 94143, USA; Neuroscience Graduate Program, University of California-San Francisco, San Francisco, CA 94143, USA

**Author notes:** Correspondence to: Kevin Yackle.

## Abstract

Opioids are perhaps the most effective analgesics in medicine. However, from 1999 to 2018, they also killed more than 400,000 people in the United States by suppressing breathing, a common side-effect known as opioid induced respiratory depression. This doubled-edged sword has inspired the dream of developing novel therapeutics that provide opioid-like analgesia without respiratory depression. One such approach has been to develop so-called ‘biased agonists’ that activate some, but not all pathways downstream of the µ-opioid receptor (MOR), the target of morphine and other opioid analgesics. This hypothesis stems from a study suggesting that MOR-mediated activation of ß2-Arrestin is the downstream signaling pathway responsible for respiratory depression, whereas inhibition of adenylyl cyclase produces analgesia. To further verify this model, which represents the motivation for the biased agonist approach, we examined respiratory behavior in mice lacking the gene for ß2-Arrestin. Contrary to previous findings, we find no correlation between ß2-Arrestin function and opioid-induced respiratory depression, suggesting that any effect of biased agonists must be mediated through an as-yet to be identified signaling mechanism.

## Introduction

More than 180 people died each day in 2018 from suppression of breathing induced by opioid overdose, a lethal effect commonly referred to as opioid induced respiratory depression (OIRD)^1,2^. Nevertheless, opioids remain among the best and most effective and widely prescribed analgesics, as evidenced by the World Health Organization’s step ladder for pain management. These two contrasting characteristics have created the urgent desire to develop or discover novel MOR agonists that provide analgesia without depressing breathing. Such a possibility emerged in 2005 when it was reported that mice with germline deletion of ß2-arrestin (*Arrb2-/-*) experience enhanced analgesia with attenuated respiratory suppression when administered systemic morphine^3^. This finding inspired a new field of ‘biased agonist’ pharmacology, which remains a central strategy in drug discovery for analgesics^4^.

Germline deletion of the µ-opioid receptor gene (*Oprm1*) completely eliminates OIRD in murine models^5^ and selective deletion of *Oprm1* within the brainstem medullary breathing rhythm generator, the preBötzinger Complex (preBötC)^6^, greatly attenuates OIRD in normoxic and hypercapnic air ^7,8^. When engaged, MOR Gi-protein signaling activates inwardly rectifying potassium channels^9^ and inhibits synaptic vesicle release^10^, thereby depressing neural signaling. Additionally, MOR can signal intracellularly in a pathway that depends on MOR internalization by *Arrb2*^11,12^. Indeed, all three pathways have been implicated in OIRD^3,13,14^ but a role for *Arrb2* has been suggested to be important and selective for respiratory depression, relative to analgesia. Since the original OIRD study of *Arrb2*-/- mice, multiple MOR agonists, dubbed “biased” agonists, have been created that signal through Gi, but not *Arrb2*, pathways, and which produce analgesia with reduced respiratory depression^15,16^. These studies have provided pharmacological support for the proposed role of ß2-Arrestin in OIRD.

More recently, however, results of these studies have been called into question^17,18^, prompting us to independently investigate the underlying necessity of *Arrb2* in mediating ORID. Here, in an experiment where we rigorously control for genetic background, we demonstrate that basal breathing and OIRD are similar in *Arrb2+/+, Arrb2*+/-, and *Arrb2*-/- littermates.

Furthermore, the *in vitro* preBötC rhythm is similarly silenced by an MOR agonist in *Arrb2*-/- and wildtype animals. Our data, together with another recent report, calls into question the necessary role of *Arrb2* in opioid induced respiratory depression and suggests that MOR biased agonists attenuate OIRD through a different mechanism.

## Results

Breathing behaviors and OIRD severity differ between strains of mice^19^. Therefore, we sought to compare basal and morphine modulated breathing in mice with the same genetic background. We bred F1 *Arrb2* +/- mice to generate littermates that were wildtype (+/+), heterozygous (+/-), and homozygous (-/-) for germline deletion of *Arrb2* (Figure 1a). Breathing was assessed by whole-body plethysmography following intraperitoneal (IP) injection of saline (for control recordings) or after IP morphine (20mg/kg, Figure 1a). This same breathing protocol was conducted in both normoxia (21% O_2_) and hypercapnia (5% CO_2_, 21% O_2_). Although hypercapnia is a non-physiologic gas, it minimizes fluctuations seen in baseline normoxia breathing rate that may confound OIRD quantification. We used our previously described analytical pipeline^7^ to assay the two key parameters that define OIRD: slow and shallow breathing. Slow breathing is measured as the instantaneous frequency of each breath and shallow breathing is defined by the peak inspiratory flow since it strongly correlates with the amount of air inspired (PIF, Figure 1b).

**Figure 1.**
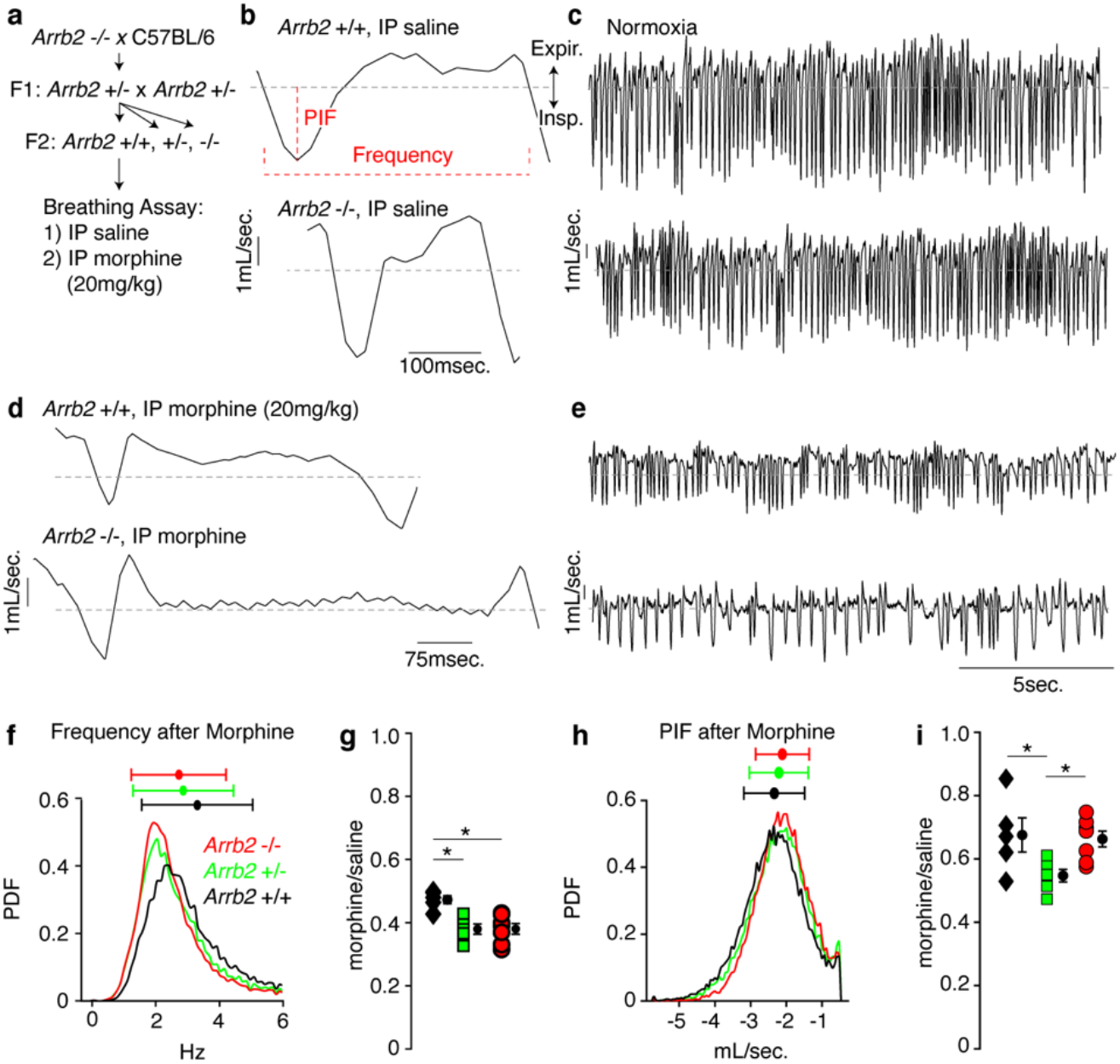
Basal respiration and OIRD in *Arrb2* littermates in normoxia. **a**, Breeding scheme to generate F2 littermates used in whole-body pleythsmography breathing recordings after intraperitoneal (IP) saline injection to establish a baseline and after IP morphine (20mg/kg). **b**, Example breath after IP saline for *Arrb2* +/+ (wildtype, top) and *Arrb2* -/- (homozygous knockout, bottom). Definition of peak inspiratory airflow (PIF, mL/sec.) and instantaneous frequency (Hz). Expir., Expiratory airflow. Insp., Inspiratory airflow. **c**, As in **b**, but example respiratory rhythm of many breaths. **d**-**e**, As in **b**-**c**, but example breaths are after IP morphine. **f**, Histogram of instantaneous respiratory frequency (Hz) for all breaths in *Arrb2*-/- (red, combined from n=7 animals), *Arrb2*+/- (green, n=6), and *Arrb2*+/+ (black, n=5) in morphine. PDF, probability density function. Top, mean (circle) ± standard deviation (bars). **g**, Ratio of average respiratory frequency in morphine to saline for *Arrb2*-/- (0.380±0.002), *Arrb2*+/- (0.379±0.002), and *Arrb2*+/+ (0.473±0.001). Mean (circle) ± standard deviation (bars). Single Factor ANOVA, F(2,15) = 9.99, P-value = 0.002. Tukey HSD post-hoc, *Arrb2*+/+ to +/- P-value = 0.004, *Arrb2*+/+ to -/- P-value = 0.003, *Arrb2*+/- to -/- P-value = 0.999. *, indicates the comparisons with P-value <0.05. Note, *Arrb2* +/+ mice have less respiratory depression, that is statistically significant, when compared to *Arrb2* +/- and -/-. **h**-**i**, As in **f**-**g**, but for peak inspiratory airflow. *Arrb2*-/- (0.663±0.004), *Arrb2*+/- (0.547±0.002), and *Arrb2*+/+ (0.676±0.014). Single Factor ANOVA, F(2,15) = 4.64, P-value = 0.027. Tukey HSD post-hoc, *Arrb2*+/+ to +/- P-value = 0.04, *Arrb2*+/+ to -/- P-value = 0.957, *Arrb2*+/- to -/- P-value = 0.050. *, indicates the comparisons with P-value <0.05. Note, *Arrb2* +/- mice have more depressed PIF, that is statistically significant, when compared to *Arrb2* +/+ and -/-. There is no statistically significant difference between *Arrb2* +/+ and -/-.

After IP saline in normoxia, the morphology of single breaths and respiratory rhythm in *Abbr2* +/+ and -/- mice appeared similar (Figure 1b-c). As expected with OIRD, after IP morphine the frequency and PIF decreased, although individual breaths and breathing rhythm remained indistinguishable in *Abbr2* +/+ versus -/- mice (Figure1 d-e). Consistently, histograms of the instantaneous frequency for each breath after IP morphine showed overlapping peaks and distributions (centered near 2Hz) in *Arrb2* +/+ (combined from n=5 mice), +/- (n=6), and -/- (n=7) animals (Figure 1f). The ratio of average instantaneous frequency after IP morphine to IP saline was similar among the genotypes, where morphine decreased respiratory rate by 60% (Figure 1g). In fact, there was a statistically significant enhanced respiratory depression in *Arrb2 -/-* and +/- mice compared to *Arrb2* +/+. Similarly, the PIF in morphine breathing histograms and ratio of IP morphine to IP saline were similar for *Arrb2* -/- and +/+ genotypes (Figure 1h-i, PIF depressed by ∼40%). These data demonstrate that OIRD in normoxia is not diminished in *Arrb2* -/- mice.

These same breathing assays and analysis were performed in hypercapnia. Hypercapnia greatly reduces the variability of breathing that exists in normoxia caused by changes in behavior, like sniffing, grooming, calmly sitting, and sleeping. If these changes differ between mice, it may confound the quantification of OIRD. Although hypercapnia increases the frequency and depth of breathing, OIRD is still observed in this condition. As anticipated, hypercapnia made breathing very regular after IP saline and IP morphine (Figure 2a-d). We observed that breath morphology, depression of frequency (depressed by ∼30%) and PIF (depressed by ∼30%) were the same among all three genotypes (Figure 2), consistent with our conclusion that OIRD is not attenuated in *Arrb2-deficient* mice.

**Figure 2.**
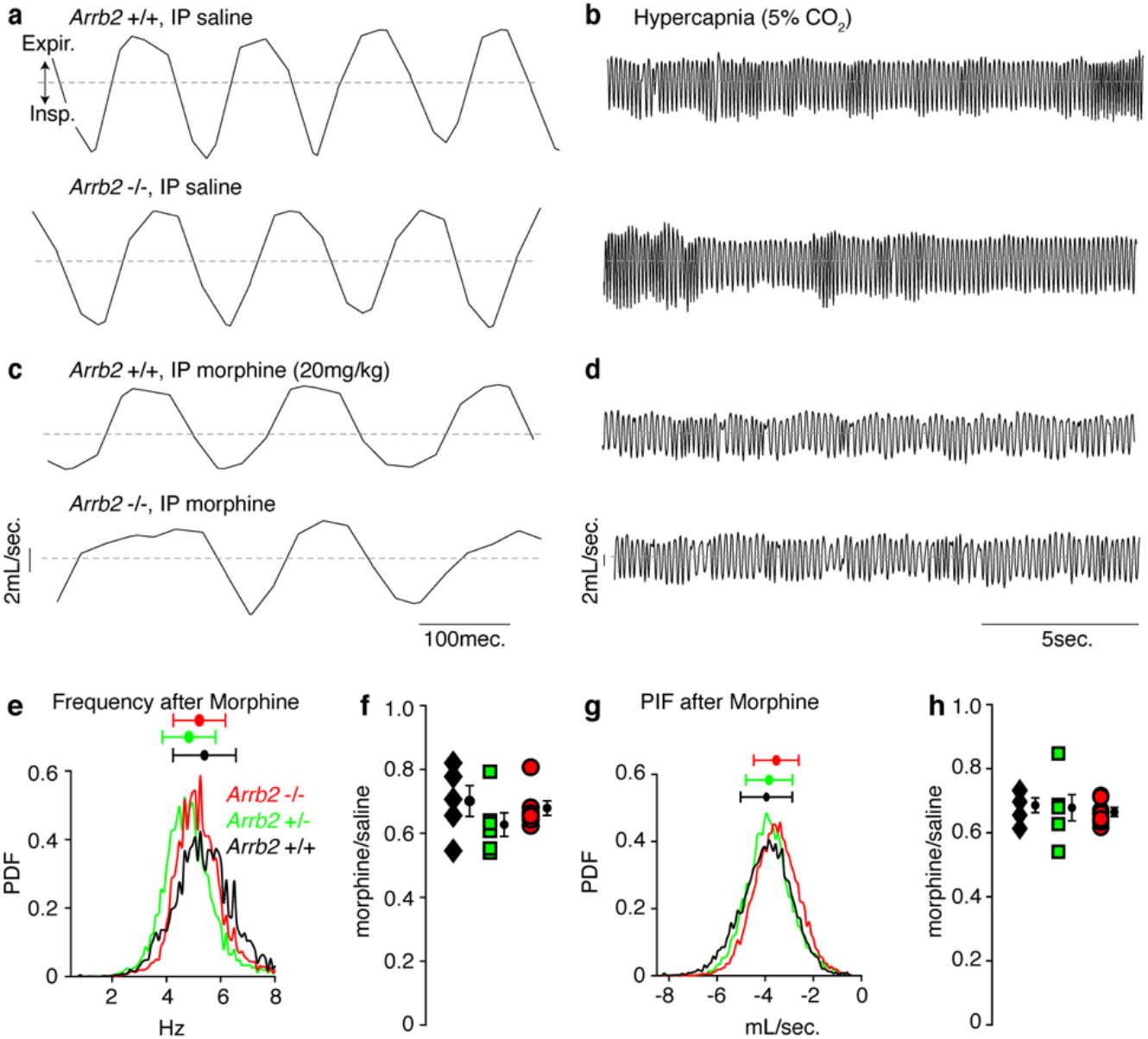
Breathing and OIRD in *Arrb2* littermates in hypercapnia. **a**-**b**, Example breaths and breathing trace after IP saline for *Arrb2* +/+ (wildtype, top) and *Arrb2* -/- (homozygous knockout, bottom) in hypercapnia (5% CO_2_, 21% O_2_). **c**-**d**, As in **a**-**b**, but after IP morphine. **e**, Histogram of instantaneous respiratory frequency (Hz) for all breaths in *Arrb2*-/- (red, combined from n=7), *Arrb2*+/- (green, n=6), and *Arrb2*+/+ (black, n=5) after IP morphine. PDF, probability density function. Top, mean (circle) ± standard deviation (bars). **f**, Ratio of average respiratory frequency in morphine to saline for *Arrb2*-/- (0.679±0.004), *Arrb2*+/- (0.627±0.008), and *Arrb2*+/+ (0.654±0.013). Mean (circle) ± standard deviation (bars). Kruskal-Wallis Test, P-value = 0.28. There is no statistically significant difference between the three genotypes. **g**-**h**, As in **g**-**h**, but for peak inspiratory airflow for *Arrb2*-/- (0.665±0.001), *Arrb2*+/- (0.678±0.010), and *Arrb2*+/+ (0.686±0.003). Single Factor ANOVA, F(2,15) = 0.15, P-value = 0.86. There is no statistically significant difference between the three genotypes.

In addition to *in vivo* breathing, we examined the effects of the opioid peptide [D-Ala, *N*-MePhe, Gly-ol]-enkephalin (DAMGO) directly on the preBötC respiratory rhythm generator and key site for OIRD. The rhythmic activity was monitored by measuring electrical activity in the hypoglossal nerve rootlet^6^. DAMGO (50nM) was bath applied to slices from *Arrb2* -/- mice, which silenced the rhythm (Figure 3, n=3). This matches the concentration of DAMGO (50-100nM) needed to silence a wildtype C57BL/6 preBötC^7^. Importantly, the rhythm was restored in the presence of the MOR antagonist, naloxone (100nM), indicating slice viability (Figure 3).

**Figure 3.**
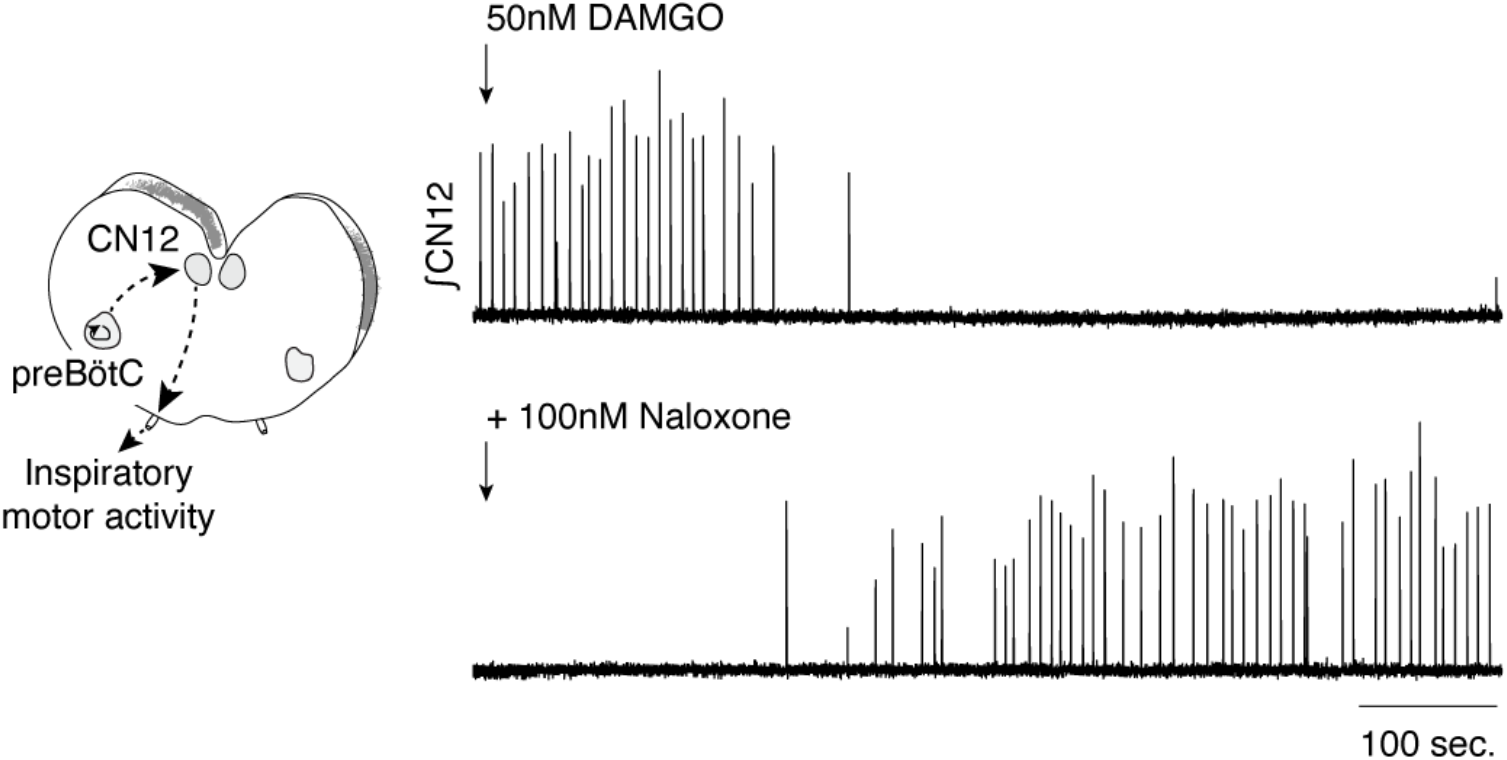
*Arrb2* -/- preBötC rhythmicity with the *Oprm1* agonist DAMGO. Left, Schematic of the neonatal medullary slice preparation containing the preBötC and the hypoglossal motor nucleus and nerve rootlet (CN12). Inspiratory bursts originating in the preBötC can be measured by suction electrodes on the CN12 rootlet (∫CN12). Right top, Bath application of 50nM DAMGO. The rhythm disappears after DAMGO is fully washed on. Bottom, Bath application of 100nM Naloxone rescues the preBötC rhythm.

## Discussion

Our *in vivo* results show that germline deletion of *Arrb2* did not attenuate opioid induced respiratory depression in normoxia nor hypercapnia. Given the effect size of previously reported results ^3^, our cohorts were sufficiently powered (n>1-4 mice minimum) to observe a decrease in OIRD if it existed. Our statistical tests were even performed conservatively to detect a negative result, meaning no difference between *Arrb2+/+* and -/-, by not correcting for multiple comparisons. Additionally, we found that the *in vitro* preBötC rhythm, a key site in OIRD^7^, is just as sensitive to DAMGO in brain slices from *Arrb2* -/- as wildtype mice. Combined, these results demonstrate that *Arrb2* germline knockout does not rescue OIRD.

This study was designed with four important features: First, we compared littermates in order to control for strain specific effects on breathing and opioid sensitivity^19^. Second, the comparison of breathing after IP injection of saline and morphine in the same animal controls for any within-animal specific breathing patterns. Third, we analyzed multiple parameters with and without averaging across breaths, in normoxia like Raehal *et al*., and also in hypercapnia to more precisely quantify OIRD. This analysis and assays should reveal even a slight rescue of OIRD.

And finally, we validated our *in vivo* studies with *in vitro* studies measuring the impact of an opioid peptide directly on the *Arrb2 -/-* preBötC.

One difference between our study and the original *Arrb2* study^3^ was our method for morphine delivery. Raehal *et al*. reported that *Arrb2* knockouts lack any respiratory depression after 20 and 50mg/kg subcutaneous morphine. In contrast, controls showed a 20 and 35% depression, respectively. However even at the maximum morphine concentration (150mg/kg) controls only showed a 40% decrease in breathing frequency whereas in our study, although we deliver only 20mg/kg morphine by IP injection, we observe a profound 60% respiratory depression in normoxia and 30% in hypercapnia in *Arrb2* +/+, +/-, and -/- mice. This respiratory depression exceeds what Raehal *et al*. reported and therefore should be sufficient to observe any OIRD rescue in *Arrb2* -/- mice.

The core premise when developing biased agonists for MOR is that *Arrb2* provides a molecular mechanism to dissociate analgesia from respiratory depression. Our findings do not support this claim. How then do some biased agonists show analgesia while minimizing respiratory depression? Perhaps biased agonists are also partial agonists. In this case, if analgesia is more sensitive than respiratory depression to the concentration of opioids, then a partial agonist might enable these two effects to be separated. This would be akin to providing a lower dose of standard opioid drugs. Regardless, our results, along with similar data from another recent study^18^, call into question the foundational model that *Arrb2* selectively mediates OIRD and suggest that in the future, we must reconsider and reinvestigate the mechanism of biased agonism *in vivo*.

## Materials and Methods

| Key Resources Table |  |  |  |  |
| --- | --- | --- | --- | --- |
| Reagent type (species) or resource |  | Source or reference | Identifiers | Additional information |
| Genetic reagent (*Mus musculus*) | *Arrb2<sup>tm1Rj</sup>/J* | The Jackson Laboratory | 011130 | RRID:IMSR_JAX:011130 |
| Genetic reagent (*Mus musculus*) | C57BL/6J | The Jackson Laboratory | 000664 | RRID:IMSR_JAX:000664 |

### Animals

*Arrb2*^20^ has been described. *Arrb2* was bred to C57BL/6 and littermates (F1) were crossed to make *Arrb2*-/-, *Arrb2*-/+, and *Arrb2*+/+ (F2). Mice were housed in a 12-hour light/dark cycle with unrestricted food and water. Mice were given anonymized identities for experimentation and data collection. All animal experiments were performed in accordance with national and institutional guidelines with standard precautions to minimize animal stress and the number of animals used in each experiment.

### Plethysmography and respiratory analysis

Plethysmography and respiratory analysis were performed as in^7^. Briefly, on the first recording day, adult (6-12 weeks) *Arrb2*-/-, *Arrb2*-/+, and *Arrb2*+/+ mice were administered IP 100µL of saline and placed in an isolated recovery cage for 15 minutes. After, individual mice were then monitored in a 450 mL whole animal plethysmography chamber at room temperature (22°C) in 21% O_2_ balanced with N_2_ (normoxia) or 21% O_2_, 5% CO_2_ balanced with N_2_ (hypercapnia). After 1 day, the same protocol was used to monitor breathing after IP injection of morphine (20mg/kg, Henry Schein 057202). Normoxia and hypercapnia morphine recordings were separated by at least 3 days. Each breath was automatically segmented based on airflow crossing zero as well as quality control metrics. Respiratory parameters (e.g. peak inspiratory flow, instantaneous frequency) for each breath, as well as averages, were then calculated. Reported airflow in mL/sec. is an approximate of true volumes. The analysis was performed with custom Matlab code available on Github with a sample dataset (https://github.com/YackleLab/Opioids-depress-breathing-through-two-small-brainstem-sites).

### Statistics

A power analysis was performed using the reported effect size from Raehal *et al*. ^3^. In this case, 1-4 mice were necessary to observe a statistically significant result. Each cohort (*Arrb2*+/+, +/-, - /-) exceeded 4. Statistical tests were performed on the ratio of IP morphine to IP saline for instantaneous respiratory frequency and peak inspiratory flow separately for normoxia and hypercapnia. A Shapiro Wilks test was first done to determine if the data was normally distributed. If normal, a single factor ANOVA was performed to determine any differences among the three genotypes (alpha <0.05). In the instance the P-value was <0.05, the Tukey HSD post-hoc test was done to determine which of the pairwise comparisons was statistically different (alpha<0.05). If the data failed to pass the Shapiro Wilks test, then the non-parametric Kruskal-Wallis test was used to determine if any differences (alpha<0.05). All the above statistics were performed using the publicly available Excel package “Real Statistics Functions” SPSS and Matlab.

### Slice electrophysiology

Rhythmic 550 to 650μm-thick transverse medullary slices which contain the preBötC and cranial nerve XII (XIIn) from neonatal *Arrb2* -/- mice (P0-5) were prepared as described ^7^. Slices were cut in ACSF containing (in mM): 124 NaCl, 3 KCl, 1.5 CaCl_2_, 1 MgSO_4_, 25 NaHCO_3_, 0.5 NaH_2_PO_4_, and 30 D-glucose, equilibrated with 95% O_2_ and 5% CO_2_ (4°C, pH=7.4). Recordings were performed in 9 mM at a temperature of 27°C. Slices equilibrated for 20 min before experiments were started. The preBötC neural activity was recorded from CNXII rootlet. Activity was recorded with a MultiClamp700A or B using pClamp9 at 10000 Hz and low/high pass filtered at 3/400Hz. After equilibration, baseline activity and then increasing concentrations of DAMGO (ab120674) were bath applied (20nM, 50nM, 100nM). After the rhythm was eliminated, 100nM Naloxone (Sigma Aldrich N7758) was bath applied to demonstrate slice viability. Rhythmic activity was normalized to the first control recording for dose response curves.

